# Regulation of light harvesting in multimeric and monomeric photosystem II supercomplexes

**DOI:** 10.1101/2020.05.04.077453

**Authors:** Eunchul Kim, Akimasa Watanabe, Christopher D. P. Duffy, Alexander V. Ruban, Jun Minagawa

**Affiliations:** Division of Environmental Photobiology, National Institute for Basic Biology, Okazaki 444-8585, Japan; Division of Biological Science, Graduate School of Science, Nagoya University, Chikusa, Nagoya 464-8602, Japan; Department of Basic Biology, School of Life Science, SOKENDAI (The Graduate University for Advanced Studies), Okazaki 444-8585, Japan; The School of Biological and Chemical Sciences, Queen Mary University of London, Mile End Road, London E1 4NS, UK

**Keywords:** photosystem II, light-harvesting complex II, photosynthesis, antenna complex, supramolecular complex

## Abstract

An intriguing architecture called ‘semi-crystalline photosystem II (PSII) array’ has been observed in the thylakoid membranes in vascular plants. It is an array of PSII–light harvesting complex II (LHCII) supercomplexes only appears in the low-light, whose functional role has not been clarified. We identified PSII–LHCII supercomplexes in their monomeric and multimeric forms in the low-light acclimated spinach leaves and prepared them using sucrose density gradient-ultracentrifugation in the presence of amphipol A8-35. When the leaves were acclimated to high-light, however, only monomeric forms were present. Single particle electron microscopy identified that the multimeric PSII–LHCII supercomplexes were composed of two (‘megacomplex’) or three (‘arraycomplex’) units of PSII–LHCII supercomplexes, which aligned like a fraction of the semi-crystalline array. Further characterization using fluorescence analysis revealed that multimeric forms have a higher light-harvesting capability, but a lower thermal dissipation capability than the monomeric form, suggesting such a configurational conversion of PSII–LHCII supercomplexes possibly serves as a structural basis for the plants’ acclimation to environmental light.

---

Light is an important energy resource for photosynthetic organisms, but its quality and quantity are variable in the field (1). Photosynthetic organisms respond and acclimate to their light conditions by controlling photosynthetic apparatus (2). In accordance with light intensities, light-harvesting, which is a primary process for photosynthesis, is optimized by modulating leave- and chloroplast-movements and tuning the number of light-harvesting complex proteins in vascular plants (3,4). Moreover, the organization of photosynthetic proteins in the thylakoid membranes are dynamically modulated according to light conditions. In vascular plants, a unique architecture, called semi-crystalline PSII array, is formed by photosystem II (PSII) and light-harvesting complex II (LHCII) supercomplexes under low-light (LL) conditions, while monomeric form of PSII–LHCII supercomplexes are predominant under high-light (HL) conditions (5,6). These configurational differences has been discussed as a potential strategy to overcome diffusion problems in crowded conditions of thylakoid membranes (7) or to modulate the capacities of non-photochemical quenching (NPQ) of the membranes (8). However, because of the technical limitation, the functional role of semi-crystalline PSII array have not been clarified.

NPQ, or qE quenching, is a photoprotection mechanism required to safely dissipate excessively absorbed light energy that tends to produce harmful reactive oxygen species (9,10). For the last few decades, the physiological importance and molecular mechanisms of qE have been intensively studied. qE is triggered by lumenal acidification of the thylakoid membranes (9,10). In vascular plants, compositions of the xanthophyll cycle pigments and PsbS are the crucial factors for qE (9). It has been proposed that a charge transfer within a chlorophyll (Chl) dimer and a zeaxanthin (Zea) or lutein (Lut) in minor LHCII proteins (11,12) and excitation transfer from a Chl cluster to a Lut in LHCII proteins (13) contributes to qE quenching. There is an alternative model that does not involve carotenoids (Car), where a charge-transfer occurs within a Chl dimer in LHCII proteins (14). These quenching processes are induced by lumenal acidification of the thylakoid membranes, which activates PsbS and violaxanthin de-epoxidase converting violaxanthin (Vio) to Zea (9,10). Both PsbS and Zea facilitate reorganization of PSII– LHCII supercomplexes and/or LHCII aggregation (10,15,16), suggesting configurational conversion of PSII–LHCII supercomplexes between a semi-crystalline array and a monomeric form described above might be linked to the modulation of NPQ (10).

Amphipols are amphipathic polymers that can be used as substitutes for detergents at the transmembrane surface of membrane proteins and, thereby, keep them soluble in detergent-free aqueous solutions (17). The most extensively studied and widely used APol, A8-35, is comprised of a mixture of short amphipathic polymers with a polyacrylate backbone onto which two kinds of side chain have been randomly grafted: ∼25% of its units carry an octylamide side chain, ∼40% an isopropylamide one, and ∼35% an underivatized carboxylate (18). Previously, we developed a procedure for purifying the PSII–LHCII supercomplex of *C. reinhardtii* employing A8-35 (19). Because the obtained complex, where the detergent dodecyl-α-D-maltoside (α-DDM) used for its solubilization were replaced by A8-35, became much more stable (19), this A8-35-substituted preparation was successfully used to determine the structure by cryo-electron microscope (20) or the binding properties of the peripheral LHCII subunits (21).

Herein, we prepared PSII–LHCII supercomplexes from spinach leaves in their monomeric and multimeric forms in the presence of A8-35. Single particle electron microscopy identified that the multimeric PSII–LHCII supercomplex, which was only identified in the LL-acclimated leaves, was composed of two or three units of PSII–LHCII supercomplexes like a fraction of the semi-crystalline array. It has a higher light-harvesting capability, but a lower thermal dissipation capability than the monomeric form. We discuss possible implications of these results with respect to the functional role of the semi-crystalline array during environmental acclimation.

## RESULTS

### Isolation of monomeric and multimeric PSII– LHCII supercomplexes using amphipol

We first isolated multimeric PSII–LHCII supercomplexes from LL-adapted spinach leaves. The thylakoid membranes were initially solubilized by α-DDM and applied to sucrose density gradient (SDG) ultracentrifugation in the presence of A8-35, which stabilized solubilized membrane protein complexes against thermodynamic dissociation (19), resulting in five green bands (Fig. 1A). Based on the analysis by SDS-PAGE (Fig. 1B), the top three bands including A1, A2 and A3 were identified as free LHC proteins, PSI–LHCI supercomplexes and PSII– LHCII supercomplexes, respectively. While these three bands correspond to the bands previously observed in the study using a green alga *C. reinhardtii*, there were two additional bands appeared in the lower part (A4 and A5) (Fig. 1A). Because the polypeptide compositions in A4 and A5 were identical as that of A3 (Fig. 1B), we presumed that A4 and A5 bands corresponded to multimer(s) of PSII–LHCII supercomplexes as reported in Albanese et al (22). Interestingly, the lower two bands (A4 and A5 bands) were not detected in the sample prepared from the HL treated leaves while the upper three bands (A1, A2 and A3) stayed the same (Fig. 1A), suggesting that the putative multimeric PSII–LHCII supercomplexes (A4 and A5) were only present in the leaves acclimated to LL conditions. The LL-dependent formation of the possible multimeric forms of PSII–LHCII supercomplexes was reminiscent to the formation of a semi-crystalline PSII array observed in LL-adapted leaves (6).

**Figure 1.**
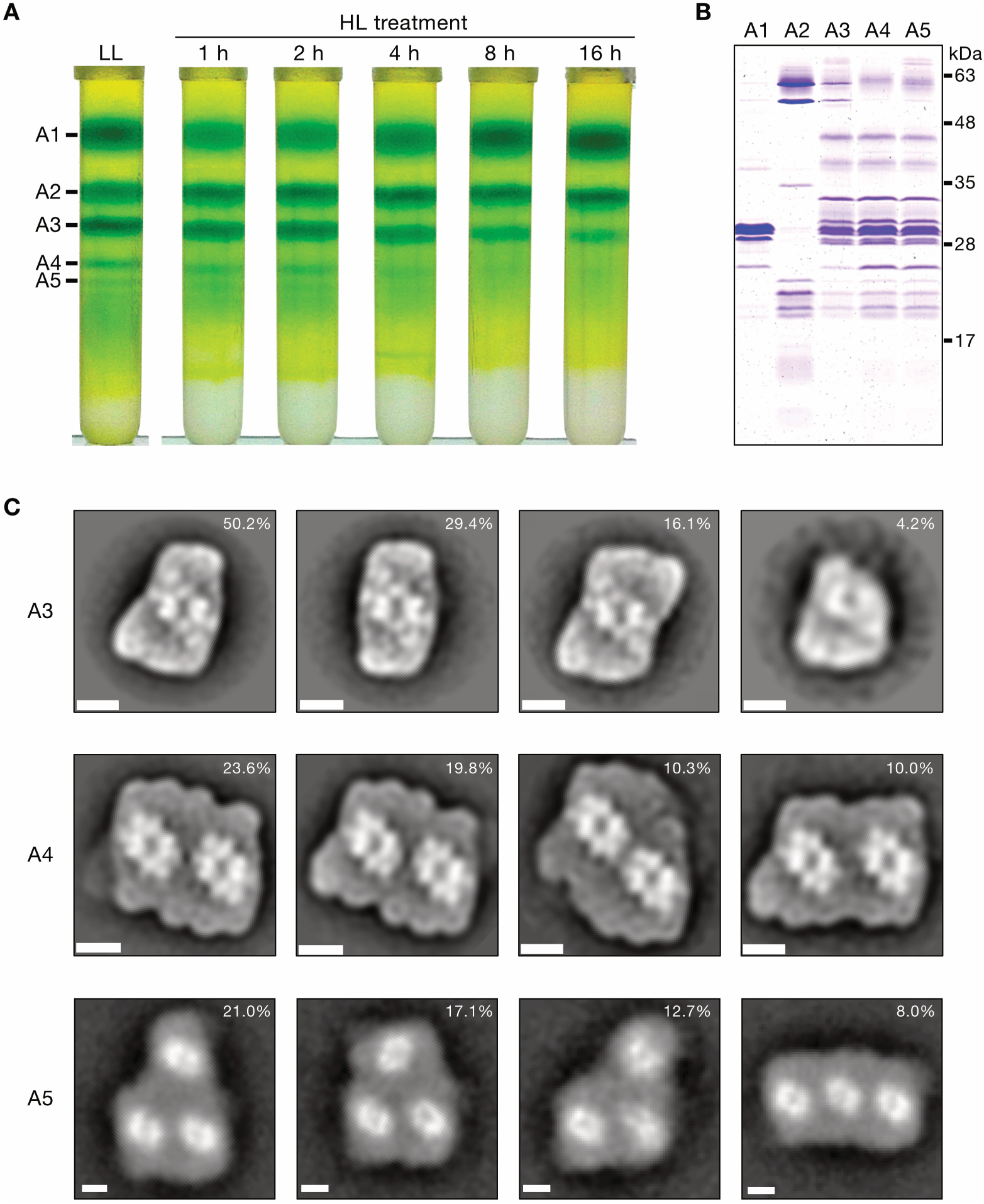
Isolation and structural characterization of photosystem complexes. **(A)** SDG ultracentrifugation of the solubilized thylakoid membranes isolated from LL-adapted leaves and after HL treatment (1000 μE m^-2^s^-1^) for 1, 2, 4, 8 and 16 hours. Thylakoid membranes isolated from spinach were solubilized with α-DDM and then replaced by amphipol A8-35 during SDG ultracentrifugation. **(B)** Polypeptides in the SDG fractions of LL-adapted leaves in (A) analyzed by SDS-PAGE stained by Coomassie brilliant blue R-250. **(C)** Averaged 2D-projection maps of negatively stained particles in A3, A4 and A5 band from the SDG fractions of LL-adapted leaves in (A). 2266, 2296 and 2626 particles, respectively for A3, A4 and A5 bands, were obtained and analyzed by three independent biological repetition. The four most abundant classes are shown, and remaining classes are shown in Fig. S2–S4. The percentages in 2D-projection maps represent the relative abundance of the particles in each band. Scale bar, 10 nm.

### Structure of monomeric and multimeric PSII– LHCII supercomplexes

PSII configurations in A3, A4 and A5 bands were characterized by single particle analysis of electron microscopy (EM) of negatively-stained particles (Fig. 1C and Fig. S1). As expected from the polypeptide compositions, A3 band was composed of monomers of PSII–LHCII supercomplexes and A4 and A5 bands were composed of multimers of PSII–LHCII supercomplexes. Four classes of monomeric PSII–LHCII supercomplexes were observed in A3 band (Fig. S2). Ten classes of PSII– LHCII supercomplex pairs were observed in A4 band (Fig. S3), which has been reported as ‘PSII– LHCII megacomplexes’ in the literature (23). Twelve classes of an array of three PSII–LHCII supercomplexes units were observed in A5 band (Fig. S4), which has never been reported before (hereafter referred to as ‘PSII–LHCII arraycomplexes’). In the other literature, similar bands with A4 and A5 bands have been isolated by using α-DDM and these bands were identified as (C_2_S_2_)_4_ and (C_2_S_2_M_2_)_2_ formed by ‘sandwiched’ PSII–LHCII supercomplexes (22,24). However, these sandwiched complexes were absent in the isolation using amphipol (20,25). In A3 band, a C_2_S_2_M_1_-type PSII–LHCII supercomplex composed of a core dimer complexes (C_2_) with two strongly bound LHCII trimers (S_2_) and one moderately bound LHCII trimer (M_1_) was dominant (50%), which was consistent with the previous report in spinach (26). The configuration of the PSII–LHCII megacomplexes in A4 band here is similar to the previously reported PSII–LHCII megacomplexes from *Arabidopsis thaliana* (23). In A5 band, PSII– LHCII arraycomplexes form various types of alignments much as like the sections in a semi-crystalline array (26) (Fig. S5), suggesting a possibility that A4 and A5 bands were fractions of a semi-crystalline PSII array.

### Pigment compositions

To characterize the optical property and pigment composition, we obtained the absorption spectrum of each band (Fig. S6A). Two main peaks around 435 and 675 nm are originated from Chl *a*, the shoulder around 650 nm is originated from Chl *b* and the shoulder around 475 nm is originated from both Chl *b* and Car. Thus, higher absorption at 475 nm but similar absorption level at 650 nm in A4 and A5 bands than that in A3 band represents the larger contributions of Car. This result is consistent with the pigment analysis using ultra-performance liquid chromatography (UPLC) (Table S1), showing A4 and A5 bands higher Car contents (Lut, Vio and Neoxanthin) per Chl *a* than A3 band. Another feature of A4 and A5 bands is that it has a red-shifted peak around 676 nm compared with A3 band. This is most likely due to the contamination of PSI because PSI shows a red-shifted absorption peak at 679.5 nm, which was supported by immunoblotting analysis of the amount of PsaA, where A4 and A5 bands contained 29% and 20% more PSI per PSII (D1) than A3 band (Fig. S7). In order to further characterize the PSII–LHCII supercomplexes, we thus adopt Chl fluorescence measurements at room temperature since most of fluorescence comes from PSII at room temperature (Fig. S6B) (27).

### Quenching properties

To assess the photoprotection capabilities, we analyzed quenching properties of A3, A4 and A5 bands by comparing Chl fluorescence decay at neutral (pH 7.5) and acidic pH (pH 5.5). NPQ_calc_ (28) was derived from fluorescence lifetimes of the non-quenching form (at pH 7.5) and the quenching form (at pH 5.5) and is used as a Stern-Volmer type NPQ parameter. As shown in Fig. 2A, the fluorescence lifetime of A3 was shorter than A4 and A5 when they were at acidic pH. NPQ_calc_ of A3 was 0.79, while those of A4 and A5 were 0.22 and 0.27, respectively (Table 1), suggesting that the monomeric PSII– LHCII supercomplexes had a much higher qE capability than the multimeric PSII–LHCII supercomplexes when in the low pH. To identify the origin of the higher qE capability in the monomeric PSII–LHCII supercomplexes, the amount of PsbS was compared because it has been reported that the PsbS could contribute qE by associating with PSII– LHCII supercomplexes (29,30). The amount of PsbS per PSII core complex (D1) was comparable in all three samples (Fig. 2B and Fig. S7), indicating that the elevated quenching capability in A3 was independent of the amount of PsbS. On the contrary, more Car was bound per Chl *a* to A4 and A5 than A3 (Table 1). A4 and A5 possessed 5% and 16% more Lut, and 13% and 24% more Vio per Chl *a* than A3, respectively. Since we analyzed the leaves adapted to LL conditions, Zea was not detected in all samples, suggesting that the elevated quenching capability of A3 here was independent of Zea. These results indicate that the higher qE capability in A3 was not due to the amount of PsbS nor Car, but could be due to conformational changes caused by monomerization of PSII–LHCII supercomplexes such as changes in the orientations of Chl(s) or Car(s). It has been actually reported that conformational changes by protein-protein interactions can activate qE process (13,31,32).

**Table 1.**
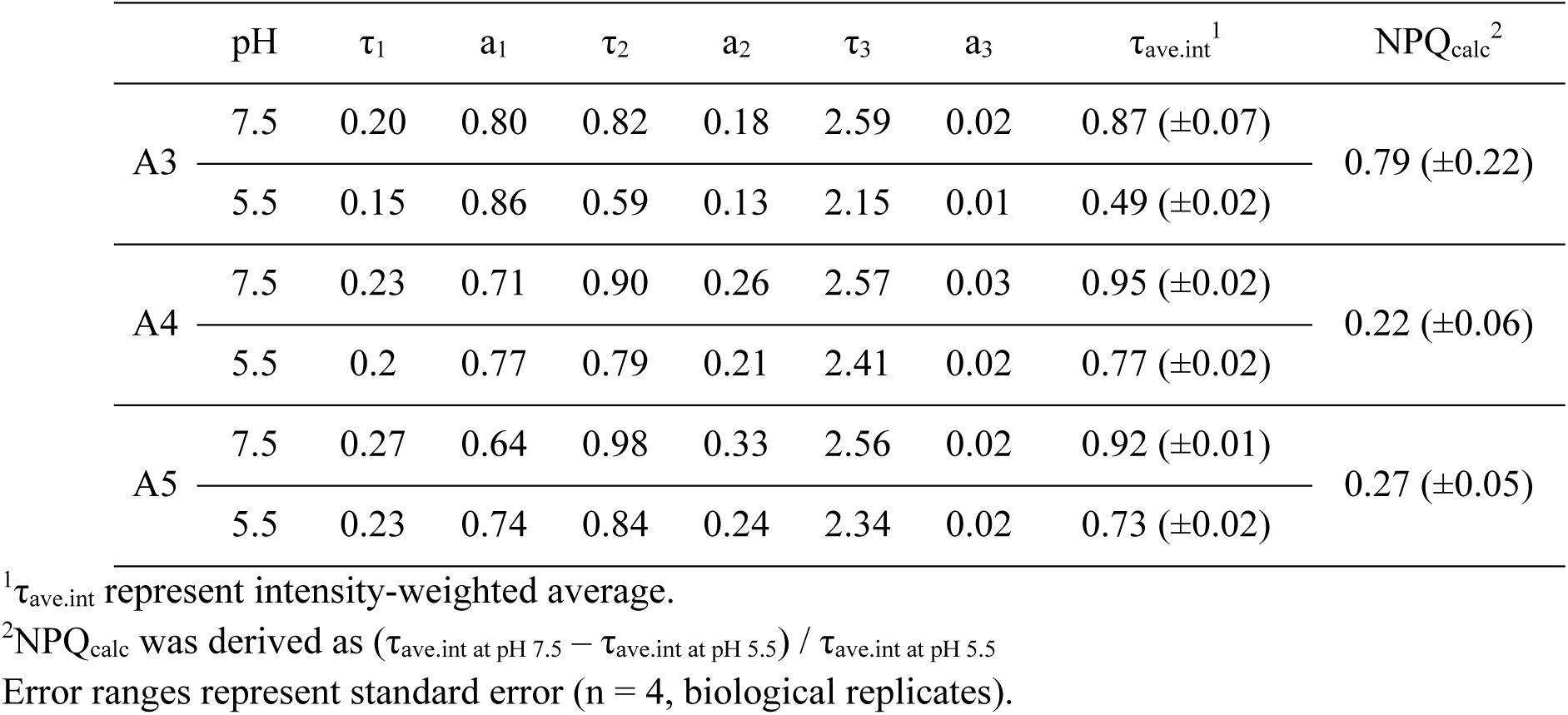
Fluorescence lifetime and pH-dependent quenching.

**Figure 2.**
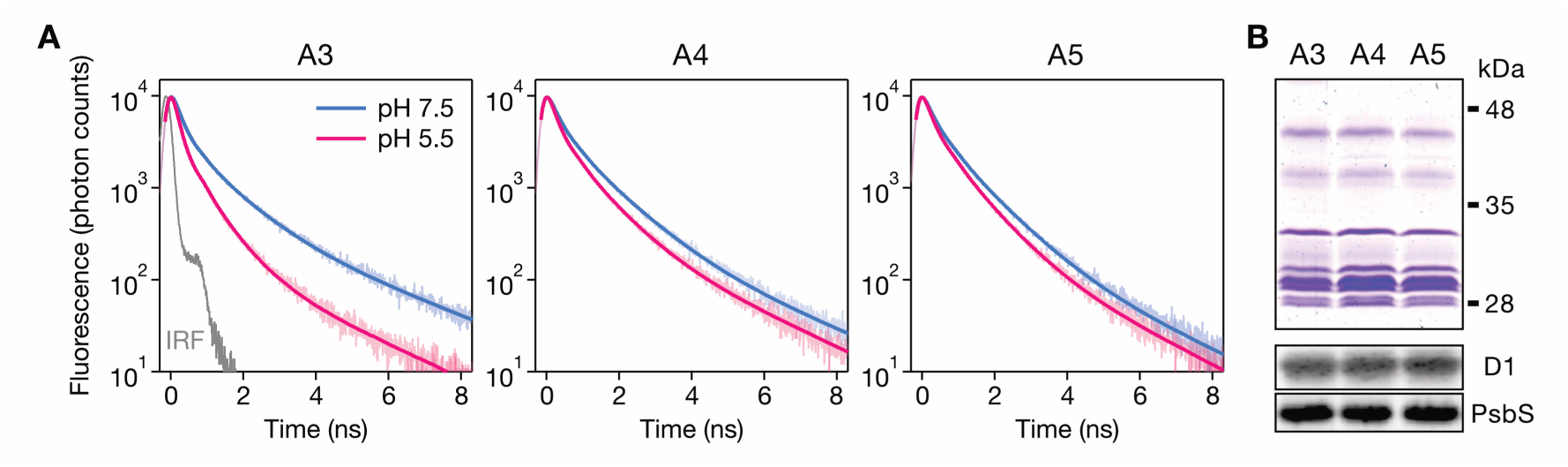
Photoprotection capability in different configurational forms of PSII–LHCII supercomplexes. **(A)** Decays of fluorescence at 685 nm obtained by excitation at 441 nm at pH 7.5 and pH 5.5. Solid lines represent fitting lines obtained by deconvolution using triple-exponential functions with instrument response function (IRF). **(B)** Polypeptides in the SDG fractions analyzed by SDS-PAGE stained by Coomassie brilliant blue R-250 and Immunoblotting analysis with antibodies against D1 and PsbS.

### Light-harvesting capabilities

The light-harvesting capabilities of monomeric and multimeric PSII–LHCII supercomplexes were investigated by fluorescence induction traces upon dark to light transition according to the previous method (33) (Fig. 3A), which determined the cross-section for PSII excitation (PSII cross-section). Because a heterogenous puddle model yielded a better fit than a homogenous puddle model or a lake model (Fig. S8), the average PSII cross-sections of the three PSII samples were estimated by the heterogenous puddle model and compared in the absence and presence of qE process (at pH 7.5 and 5.5, respectively). At pH 7.5, A4 and A5 showed 10.0% and 37.6% larger PSII cross-sections than A3, respectively (Fig. 3B). At pH 5.5, however, A4 and A5 showed 53.3% and 80.9% larger PSII cross-sections than A3, respectively (Fig. 3B). Multimerization therefore contributes to increase the PSII cross-section of the PSII–LHCII supercomplexes.

**Figure 3.**
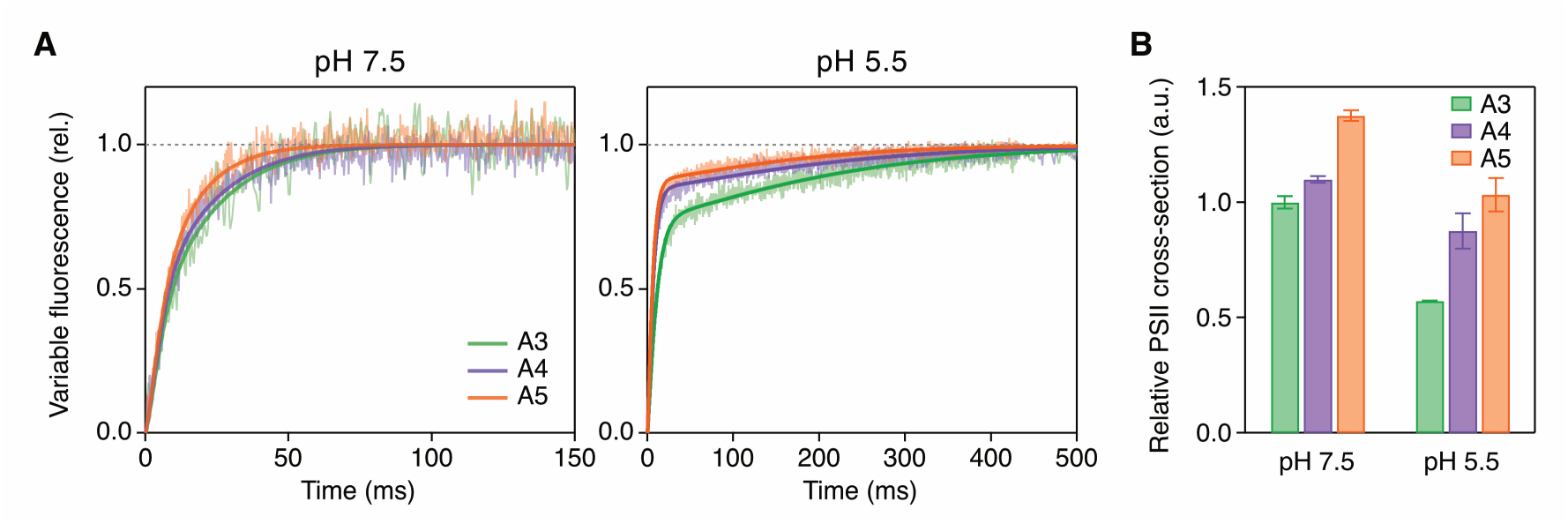
Light-harvesting capabilities in different configurational forms of PSII–LHCII supercomplexes. **(A)** Fluorescence induction traces upon dark to light transition (28 μE m^-2^s^-1^) at pH 7.5 and pH 5.5 in the presence of 20 μM DCMU. Each trace represents the averaged trace of three independent measurement. Solid lines represent the simulated curves by heterogenous puddle models. **(B)** Relative PSII cross-section at pH 7.5 and pH 5.5. Error bars represent 95% confidence intervals of obtained model parameters.

We then examined whether the larger PSII cross-sections in A4 and A5 were due to larger physical antenna size or higher light-harvesting efficiency (energy transfer efficiency from antenna to the core), or both. The physical antenna sizes of the PSII– LHCII supercomplexes in each sample were evaluated based on the result of single particle analysis by EM (Fig. S5–S7 and Table S2 and S3). The estimated physical antenna sizes of PSII–LHCII supercomplexes in A4 and A5 were 9% and 14% larger than that in A3 (Table 2). This result is consistent with the lower Chl *a*/*b* ratio in A4 and A5 (Table S1) which represents more antenna proteins, even though the PSI contamination in A4 and A5 bands could increase Chl *a*/*b* ratio. The light-harvesting efficiencies were deduced by ratios of PSII cross-sections to physical antenna sizes. The results indicate that A4 and A5 have 41% and 59% higher light-harvesting efficiencies than A3 at pH 5.5, respectively, while A4 and A5 have 1% and 21% higher light-harvesting efficiencies than A3 at pH 7.5 (Table 2). Therefore, the multimeric formation of PSII–LHCII supercomplexes increase light-harvesting efficiency as well as physical antenna sizes associating more antenna proteins. Interestingly, their light-harvesting efficiencies were greatly reduced by monomerization at the acidic pH, which was correlated with the increased level of NPQ capacities, suggesting that the energy transfer from antenna to the core could be inhibited in the monomerized forms, but not in the multimeric forms of the PSII–LHCII supercomplexes.

**Table 2.** Physical antenna size and light-harvesting efficiency.

|  | The number of Chls per PSII core dimer<sup>1</sup> | Relative physical antenna size | Relative light-harvesting efficiency (a.u.)<sup>2</sup> |  |
| --- | --- | --- | --- | --- |
|  |  |  | at pH 7.5 | at pH 5.5 |
| A3 | 251 ($\pm 37$) | 1.00 | 1.00 ($\pm 0.03$) | 0.57 ($\pm 0.20$) |
| A4 | 274 ($\pm 20$) | 1.09 | 1.01 ($\pm 0.02$) | 0.80 ($\pm 0.09$) |
| A5 | 285 ($\pm 28$) | 1.14 | 1.21 ($\pm 0.03$) | 0.91 ($\pm 0.08$) |

<sup>1</sup>The number of Chls per core dimer was estimated based on the single particle analysis result. All values are mean ± SD of analyzed particles (2266, 2296 and 2626, respectively) obtained by three independent experiments in Fig. S2–S4 and Table S3. The relative physical antenna size was estimated from the average number of Chls per PSII core dimer.
<sup>2</sup>Relative light-harvesting efficiency was calculated by dividing PSII cross-section by the relative physical antenna size. Error ranges represent 95% confidence intervals of obtained model parameters for estimating PSII cross-section.

## DISCUSSION

In this study, using a recently established amphipol A8-35-employed protocol (19,21), we isolated monomeric and multimeric PSII–LHCII supercomplexes, putative fractions of semi-crystalline PSII arrays (Fig. 1). Fluorescence analysis combined with single particle analysis revealed that the multimeric PSII–LHCII supercomplexes have higher light-harvesting capability but lower photoprotection (qE) capability than the monomeric PSII–LHCII supercomplexes (Fig. 2 and 3). These results suggest a possibility that the segregation of semi-crystalline PSII arrays to monomeric PSII–LHCII supercomplexes decreases their light-harvesting capability but increases their photoprotection capability.

Based on these results, we propose a configurational modulation model of light-harvesting properties of PSII–LHCII supercomplexes (Fig. 4). Under LL conditions, PSII– LHCII supercomplexes are in the semi-crystalline state, which has an advantage in light-harvesting (Fig. 4A). During acclimating to HL conditions, reorganization of PSII–LHCII supercomplexes occur in the membrane, converting a semi-crystalline PSII array into a disordered assembly of the PSII–LHCII supercomplexes (Fig. 4B). Some of the tightly associated LHCIIs become peripheral during this disordering process. The reduced physical antenna size of the monomeric PSII–LHCII supercomplexes (Table 2) implies some of the LHCIIs dissociate from PSII–LHCII supercomplexes during biochemical treatment of the membranes in the disordered state. The dissociation of LHCIIs under HL conditions is consistent with the previous observations (10,16), which has been implicated as a critical step for the aggregation of LHCIIs required for qE as well as the contributions of PsbS and Zea (10,15). Membranes with such a disordered PSII–LHCII supercomplexes could have a higher qE capability, where charge-transfers between Zea and Chls within CP29 (11,12), excitation-transfers from Chl to Lut within LHCII (13), etc., possibly occur at the acidic pH. This sort of configurational modulation enables adjustment of qE capability according to the environmental light conditions, which eventually contributes photosynthesis yield (34).

**Figure 4.**
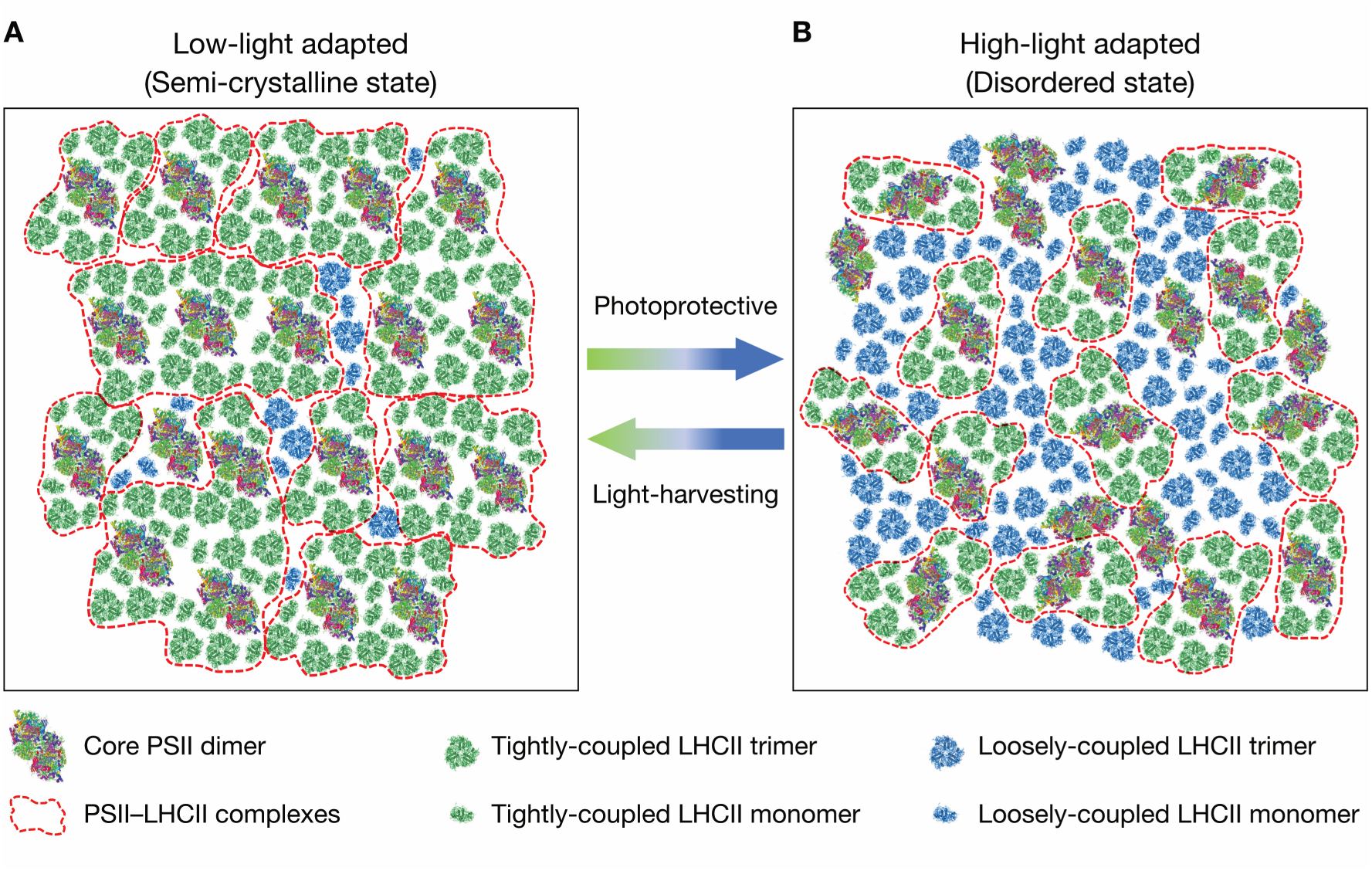
A proposed model for photoacclimation of PSII–LHCII supercomplexes. Schematic representation of the putative arrangements of PSII–LHCII supercomplexes in the thylakoid membrane in LL **(A)** and in HL **(B). (A)** 20 PSII complexes are in the semi-crystalline array state. **(B)** 20 PSII complexes are in the disordered state. In this state, LHCIIs tightly and loosely coupled to PSII have a potential to exert NPQ upon lumenal acidification (photoprotection capability). The complexes outlined by *dashed red lines* could be isolated after detergent solubilization.

The other important issue arisen from this study is the factor triggering the configurational modulation of PSII–LHCII supercomplexes. We hypothesize four factors: (i) acidic pH, (ii) phosphorylation of LHC and/or PSII core, (iii) the activation of PsbS, (iv) accumulation of Zea. (i) Our previous study (21) showed that the acidic pH condition induces conformational change of complexes so that binding of S-trimer to the PSII core become loose. We suppose that such HL-adapted condition induces sufficiently long acidify condition which can induce the conformational change. (ii) It has been well studied that the STN7 and STN8-dependent phosphorylation induces detachment of LHCII trimers from PSII and reorganization of photosynthetic protein complexes (35,36). We suppose such a large reorganization can induce configurational modulation of PSII–LHCII supercomplexes. (iii) It has been shown that PsbS regulates the membrane fluidity, which is an important factor for reorganization of photosynthetic protein complexes (15). Therefore, it is plausible that the activation of PsbS induces the configurational modulation of PSII–LHCII supercomplexes (8). (iv) It is well known that the HL condition induces de-epoxidation of Vio (37,38). We suppose that change of xanthophyll species can induces the configurational modulation of PSII–LHCII supercomplexes due to its different chemical structure and property. These hypotheses can be verified by further study using several mutants lacking PsbS, STN7 or STN8 and xanthophyll cycle.

Our findings provide a feasible explanation for the discovery of Goral et al. (39) that showed a correlation between the semi-crystalline PSII array *in vivo* and the reduced NPQ in LHCII and PsbS mutants in *Arabidopsis*. This could be at least partially explained by the reduced mobility of the PSII components and restricted freedom for the rearrangement that is required for a transition into the NPQ state, when in the crystalline arrangement. The details of the molecular mechanism of how this configurational change occurs remains to be investigated by structural and ultrafast spectroscopic studies.

## EXPERIMENTAL PROCEDURES

### Isolation of Thylakoid Membranes and Photosystem Complexes

Spinach leaves were quickly frozen in liquid nitrogen after light treatments and disrupted using a blender with a buffer containing 0.33 M sucrose, 5 mM MgCl_2_, 1.5 mM NaCl and 25 mM MES (pH 6.5, NaOH). From the suspension obtained by blending, thylakoid membranes were isolated according to the same protocol used for *C. reinhardtii* (40), and were suspended in a 25 mM MES buffer (pH 6.5) containing 1 M betaine at 0.3 mg Chl/mL and solubilized with 1.0% α-DDM (Anatrace, Maumee, OH) for 10 min on ice. Unsolubilized thylakoids were removed by centrifugation at 25000 *g* for 1 min. After centrifugation, A8-35 (Anatrace) was added at a final concentration of 1% to the solubilized thylakoids for amphipol substitution, and subjected to sucrose density-gradient (SDG) ultracentrifugation as previously described (19). For spectroscopic analysis of isolated PSII–LHCII complexes, samples were diluted to 2 μg of Chl per mL by a 25 mM MES buffer (pH 6.5) containing 1 M betaine.

### SDS/PAGE and Immunoblotting Analysis

SDS/PAGE and immunoblot analyses were performed as previously described (41). For Coomassie Brilliant Blue R-250 staining, 1.0 μg Chl of samples were applied to SDS/PAGE. For immunoblot analyses, 0.3 μg Chl of samples were applied to SDS/PAGE. Antibodies for D1 and PsbS proteins were purchased from AgriSera (AS05-084 for D1, AS09-533 for PsbS). The antibody used for PsaA detection was described previously (41). An anti-rabbit horseradish peroxidase-conjugated antiserum (#7074, Cell Signaling Technology, Danvers, MA, USA) was used as a secondary antibody. Densitometric analyses of the detected images were performed using Image Lab software (Bio-Rad).

### Electron Microscopy and Single-particle Analysis

The isolated complexes were diluted to 2 μg of Chl per mL in the 25 mM MES buffer, applied to glow-discharged carbon-coated copper grids, and negatively stained for 30 s with 2% uranyl acetate three times. Electron micrographs were obtained using a JEM 1010 electron microscope (JEOL; Japan) at 80 kV and 150,000× magnification. Images were recorded using an Olympus Velete CCD camera (2,048 × 2,048) at a pixel size of 5.0 Å. In total, 100 micrographs for PSII–LHCII supercomplexes, 400 micrographs for PSII–LHCII megacomplexes, 600 micrographs for PSII–LHCII arraycomplexes were collected. The Relion 2.1 package was used for 2D classification (42).

### Pigment Analysis

Pigments of PSII supercomplexes extracted by 80% acetone and analyzed by ultra-performance liquid chromatography (UPLC) using a H-class system (Waters) as previously described (43).

### Time-resolved fluorescence decay analysis

Time-resolved fluorescence decays were obtained by a time-correlated single photon counting system (FluoroCube; HORIBA Jobin-Yvon) at room temperature. Samples were excited at 441 nm with a repetition rate of 1 MHz by a diode laser (N-440L; HORIBA Jobin-Yvon) and emissions were detected at 685 nm with 32 nm bandwidths.

### Fluorescence induction traces measurements and analysis

Prior to the measurement of fast fluorescence induction traces, PSII–LHCII complexes were kept in dark for 6 h for fully oxidizing PSII reaction centers (Fo). Fast fluorescence induction traces, from Fo (open reaction centers) to Fm (closed reaction centers), were measured under weak light conditions (28 μE m^-2^s^-1^) in the presence of 20 μM of 3-(3,4-dichlorophenyl)-1,1-dimethylurea (DCMU). The sample concentrations were set to 10 μg of Chl per mL. The concentration of DCMU has been set as enough concentration for saturated conditions (33).

To simulate the fast fluorescence induction of the samples we used the procedure previously outlined in the Supplementary Information of a previous study (33), which was based on a relevant mathematical model (44). Here, as in the previous study, it was found that a satisfactory fit is only obtained for a heterogeneous puddle model. Indeed the ‘lake’ model can be discounted by simple visual inspection of the traces due to their obvious lack of sigmoidicity. In the ‘puddle’ model each antenna puddle contains a finite (and small) number of reaction centers (RCs), *N*. A full derivation of the overall theory can be found in the previous study (33). Here we briefly summarize the key equations. The normalized fluorescence induction trace, in the presence of qE quenchers, is defined,

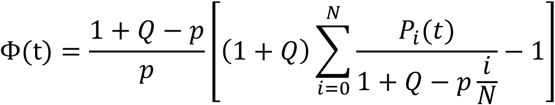

where,

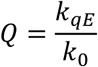

Is the ratio between the rate constants of excitation trapping by the qE quenchers, *k*_*qE*_, and excitation trapping by open RCs, *k*_0_. Similarly,

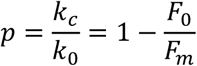

Describes (background) quenching in conditions of closed reaction centers. The summation index *i* counts the number of *closed* RCs. For each value of *i* we assign a time-dependent probability, *P*_*i*_(*t*). These probabilities evolve according to the Master Equations,

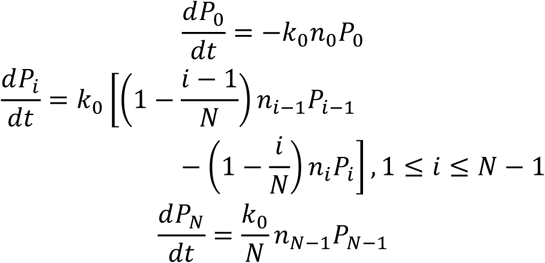

Where *n*_*i*_ is the steady state excitation density within a system with *i* closed RCs,

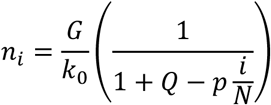

*G* is the rate constant for excitation generation in the antenna assuming constant light intensity, *I*. It is directly proportional to the cross section, *σ*, of the antenna,

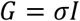

As *N* increases lake-type (sigmoidal) behavior is recovered. The best fits were obtained for *N* = 2 (i.e. a PSII RC dimer) in all cases. Moreover, the traces are visibly biphasic and so good fits could only be obtained if we assumed heterogeneity. For simplicity we assumed two sub-populations. The fluorescence trace is then given by,

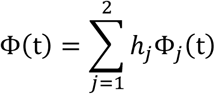

Where,

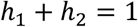

In total our fitting parameters are *G*_1_,*G*_2_, *h*_1_, *Q*_1_ and *Q*_2_. The parameter *p* is obtained directly from the data and is assumed to be identical for each subpopulation and independent of pH. To avoid over-fitting we adopt a careful procedure. First we fit the traces for pH 7.5 and assume that *Q*_1_ = *Q*_2_ = 0. Any residual quenching present is quantified by *p*. Next, the long-time kinetics (i.e. as the curve approaches the maximal fluorescence yield) are defined almost entirely by the sub-population with the smallest cross-section which we define as *G*_1_.We therefore obtain *G*_1_ by fitting the long time data to,

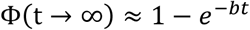

Where,

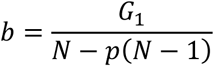

*G*_2_ and *h*_1_ are then obtained by fitting the full data with fixed *G*_1_. At pH 5.5 we assume the presence of qE quenchers, *Q*_1_ and *Q*_2_, Additionally, we allow for changes in the cross-sections, *G*_*j*_, of the two sub-populations but we use their pH 7.5 values as initial guesses. For simplicity we assume that *h*_1_ does not change.

To find an appropriate model for estimating PSII cross-section of the three PSII–LHCII supercomplexes, we first compared fitting results derived from ‘puddle’ model with those from ‘lake’ model. For all three samples, a puddle model with two reaction centers showed a better fit rather than a ‘lake’ model (Fig. S8), implying that the PSII–LHCII supercomplexes forming mega- or arraycomplexes are not tightly coupled for energy transfer. Even within the array complexes the each dimeric PSII core appears to be functionally separated in terms of energy transfer.

## Author contributions

E.K., A.W. and J.M. designed research. A.W. performed biochemical experiments and electron-micrograph analysis. E.K. performed spectroscopic experiments and analyzed data. C.D.P.D, E.K., A.V.R, J.M. analyzed antenna cross-section. E.K. and J.M. wrote the manuscript. All authors reviewed and approved the final version of the manuscript.

## Acknowledgements

We thank Dr. Ryutaro Tokutsu for helpful comments and Dr. Shunichi Takahashi for providing *Arabidopsis thaliana*. We also thank to Chiyo Noda for assisting in pigment composition analysis.

## FOOTNOTES

This work was supported by Japan Society for the Promotion of Science (JSPS) KAKENHI Grant-in-Aid 16H06553 (to JM).

